# Melt electrowritten scaffolds with bone-inspired fibrous and mineral architectures to enhance BMP2 delivery and human MSC osteogenesis

**DOI:** 10.1101/734855

**Authors:** Kian F. Eichholz, David A. Hoey

**Affiliations:** Dept. Mechanical, Aeronautical and Biomedical Engineering, Materials and Surface Science Institute, University of Limerick, Limerick, Ireland; Trinity Centre for Bioengineering, Trinity Biomedical Sciences Institute, Trinity College Dublin, Ireland; Dept. of Mechanical and Manufacturing Engineering, School of Engineering, Trinity College Dublin, Ireland; Advanced Materials and Bioengineering Research Centre, Trinity College Dublin & RCSI

## Abstract

Material micro-architecture and chemistry play pivotal roles in driving cell behaviour. Bone at a cellular level consists of arranged fibres with a cross-fibrillar mineral phase made up of curved nano-sized needle shaped crystals. This nano-structured mineral architecture can bind and stabilise proteins within bone for centuries and thus holds promise as a strategy for therapeutic delivery in regenerative medicine. Herein, we use melt electrowriting (MEW) technology to create fibrous 3D PCL micro-architectures. These scaffolds were further modified with an extrafibrillar coating of plate shaped micron-sized calcium phosphate crystals (pHA), or with a novel extrafibrillar coating of needle shaped nano-sized crystals (nnHA). A third scaffold was developed whereby nano-sized crystals were placed intrafibrillarly during the MEW process (iHA). X-ray diffraction revealed altered crystal structure and crystallinity between groups, with hydroxyapatite (HA) being the primary phase in all modifications. Water contact angle was investigated revealing increased hydrophilicity with extrafibrillar coatings, while tensile testing revealed enhanced stiffness in scaffolds fabricated with intrafibrillar HA. Biological characterisation demonstrated significantly enhanced human stem/stromal cell mineralisation with extrafibrillar coatings, with a 5-fold increase in mineral deposition with plate like structures and a 14-fold increase with a needle topography, demonstrating the importance of bone mimetic architectures. Given the protein stabilising properties of mineral, these materials were further functionalised with BMP2. Extrafibrillar coatings of nano-needles facilitated a controlled release of BMP2 from the scaffold which further enhanced mineral deposition by osteoprogenitors. This study thus outlines a method for fabricating scaffolds with precise fibrous micro-architectures and bone mimetic nano-needle HA extrafibrillar coatings which significantly enhance mesenchymal stem/stromal cell (MSC) osteogenesis and therapeutic delivery and thus hold great promise for bone tissue regeneration.

## 2 Introduction

Autografts are the gold standard for bone defect repair due to their osteoconductive, osteoinductive and immunocompatible properties, but unfortunately are limited in supply [1]. Due to technology advancements and an improved understanding of the physiological mechanisms of bone repair, synthetic tissue engineering approaches have demonstrated great progress in developing bioinspired materials that can promote bone formation [2]. These approaches continue to fall short of the gold standard however, therefore, continued development is required. Bone is a complex, hierarchical multi-phasic organ comprised primarily of mineralised collagen fibrils spanned by additional continuous, cross-fibrillar mineral units [3, 4], It was only recently identified that the first fundamental component in this hierarchy is composed of needle-shaped mineral units of base 5 nm and length 50 - 100 nm, which merge to form platelets and mineral stacks and aggregate to form irregular 3D structures of size 200 - 300 nm which are crossfibrillar in nature [4]. It is the combination of collagen and mineral with specific topographies, working in unison in this hierarchical structure, which gives the bone both its excellent biological and mechanical properties. Moreover, this mineral nano-topography identified in bone is believed to be a key contributor to non-collagenous protein binding and stabilisation, preserving proteins for centuries within the tissue without denaturation [5]. This therefore provides a potential strategy for therapeutic delivery in regenerative medicine [6]. Recapitulating this unique multiscale architecture may represent an innovative approach to develop effective bioinspired materials to promote bone repair.

Bone formation occurs following mesenchymal stem/stromal cell osteogenic differentiation into an osteoblast, which begins laying down collagen fibrils and enriching the surrounding environment with alkaline phosphatase and osteocalcin to promote mineral deposition within the collagen template [7, 8]. The resulting nano-crystals are initially formed within collagen fibrils, eventually nucleating to fill the voids forming a composite structure of mineral and collagen fibres approximately 10 μm in diameter [9]. The underlying collagen fibrillar template is thus crucial in dictating cellular behaviour in bone and many other tissues, with dysregulation of this unique structure leading to undesirable outcomes including cell death, abnormal differentiation and tumour development [10, 11]. Through the advent of new technology such as 3D printing, precise control of scaffold architecture is now achievable [12]. Fused deposition modelling (FDM) can produce scaffolds with defined fibre architecture creating pores of differing shapes [13] in addition to facilitating mineral incorporation [14–18], which in turn drives mesenchymal stem/stromal cell (MSC) differentiation down specific lineages. However, FDM is limited in that fibre diameters below 100 μm are difficult to fabricate. These relatively large fibres result in a pseudo-3D environment at the scale of the cell [19]. Electrospinning is a common fabrication technique to produce physiological fibre diameters on the nano/micron scale which have been utilised in bone tissue engineering strategies [20, 21]. However, due to the unstable "whipping" motion of the electrospinning jet during fibre deposition, there is limited control over fibre architecture using this approach. Melt Electrospinning Writing (MEW) is a recently developed 3D printing technology which overcomes the above limitations of FDM and electrospinning, being capable of fabricating and controlling the deposition of micron-scale fibres [22–24]. This facilitates the manufacture of complex fibrous micro-architectures [25–27] consistent with that seen in native extracellular matrix (ECM) including that of bone. We recently utilised MEW to fabricate scaffolds with bone inspired fibre diameters of 10 μm and investigated whether the fibre architecture in terms of degrees of alignment could influence stem/stromal cell behaviour [28]. Interestingly, the combination of fibre diameter and alignment had significant impact on cell shape, cell signalling and ultimately MSC osteogenesis, with a 90° fibre architecture mediating a 4-fold increase in stem/stromal cell mineralization when compared to 10° aligned architectures or a traditional random electrospun scaffold, demonstrating the critical role of fibrillar architecture in mediating stem/stromal cell behaviour and highlighting MEW as a powerful tool to fabricate physiological bioinspired environments for tissue engineering strategies.

In bone tissue, the mineral component makes up 60-70% of its dry weight, and is composed primarily of non-stoichiometric apatite along with other loosely bound ions including Ca^2+^, HPO_4_^2-^, C0_3_^2-^ and Mg^2+^ in a hydrated layer at the surface [29]. As with fibrillar architecture, the mineral structure in bone plays a key role in mediating cell behaviour, with deficiencies or changes in mineral composition contributing to diseases such as osteoporosis [30]. This has been exploited in a host of strategies for orthopaedic and dental applications, where materials and surfaces have been modified with calcium phosphates with the aim of creating biomimetic materials which promote osseointegration and tissue regeneration. This includes coatings for implant fixation, cements and adhesives for orthopaedic and dental implants, and synthetic bone grafts [31]. A wide range of commercially available materials have been investigated, including hydroxyapatite (HA), calcium deficient apatite (CDA), α-and β tricalcium phosphate (TCP) and biphasic calcium phosphates such as HA + TCP, with a range of other materials including dicalcium phosphate (DCP) and Tetracalcium phosphate (TTCP) also being investigated [32]. As with fibrillar architecture, an ever increasing appreciation for the importance of the architecture and particle size of the mineral phase is being gained, which defines surface topography and thus has a considerable influence on cell behaviour and scaffold performance [33]. When cultured on nano-scale HA (nHA) films with varying particles sizes from 20 - 80 nm in diameter, MSCs exhibit a trend of greater viability and proliferation with smaller particle sizes [34]. The osteogenic capacity of nHA has been further demonstrated in a range of studies where cells have been cultured in nHA suspensions with particles of size between 20 - 100 nm, with cells demonstrating uptake of particles and exhibiting enhanced osteogenic differentiation [35–37]. The role of HA particle size has also been investigated in the 3D printing of composite PCL-HA blends, with enhanced proliferation and alkaline phosphatase (ALP) activity of MSCs seen with nano-particles compared to micro-particles [38]. Remarkably, particle size has not only been shown to be an important cue for MSC osteogenesis, but also plays a role in suppressing undesired cellular outcomes in bone. This has been demonstrated via osteosarcoma inhibition [34] and reduced melanoma proliferation [39] with nHA compared to larger particles, further reinforcing the therapeutic potential of nano-scale particles for additional applications such as bone regeneration following tumour ablation. A recent study has also demonstrated the regenerative capacity of grafts with a needle surface topography *in-vivo*, with sub-micron needles triggering earlier and accelerated bone formation [40].

Bone tissue also houses non-collagenous proteins and growth factors embedded within its composite matrix, which are remarkably resolute and can be preserved for centuries due to the great stability provided by the mineral structure [5]. It is thus of no surprise that the great binding affinity to HA of a host of biological and synthetic components has been exploited for therapeutic benefits. HA has been extensively used as a delivery mechanism for BMP2 protein with proven success in a number of in-vivo models [41–43], while nHA particles have also been used for non-viral gene delivery of BMP2 to enhance MSC osteogenesis [44, 45]. Anti-tumour drugs have also been combined with nHA as a delivery vector, with potential applications in osteosarcoma chemotherapy [46] and the development of targeted drug delivery methods [47]. Interestingly, protein binding and stabilization is closely related to the nanoscale structural features of the mineral coatings [6], demonstrating the importance of controlling this nHA architecture. In addition to the already pro-osteogenic capabilities of biomimetic nHA, there is great potential to exploit its natural affinity to further advance treatments by facilitating therapeutic delivery providing an extensive range of enhanced regenerative benefits.

In this study, we aim to recapitulate the native structure of bone via the development of a biomimetic tissue regeneration strategy comprised of a micro-fibrous matrix spanned by an intrafibrillar nano-needle mineral phase stabilising bound protein for enhanced tissue regeneration. Herein, we use melt electrowriting (MEW) technology to create fibrous 3D PCL micro-architectures. These scaffolds were further modified with an extrafibrillar coating of plate shaped micron-sized calcium phosphate crystals (pHA), or with a novel extrafibrillar coating of needle shaped nano-sized crystals (nnHA). A third scaffold was developed whereby nano-sized crystals were placed intrafibrillarly during the MEW process (iHA). Along with vastly differing surface topographies, surface chemistry was also altered, with calcium/phosphate ratios within the range seen in healthy native bone. Biological characterisation demonstrated significantly enhanced human stem/stromal cell mineralisation with extrafibrillar coatings, with a needle topography producing the most robust bone formation, demonstrating the importance of bone mimetic architectures. Given the protein stabilising properties of mineral, these materials were further functionalised with BMP2. Extrafibrillar coatings of nano-needles facilitated a controlled release of BMP2 from the scaffold which further enhanced mineral deposition by osteoprogenitors. This study thus outlines a method for fabricating scaffolds with precise fibrous micro-architectures and bone mimetic nano-needle HA extrafibrillar coatings which significantly enhance MSC osteogenesis and therapeutic delivery and thus holds great promise for bone tissue regeneration.

## 3 Materials and methods

### 3.1 Scaffold fabrication

#### 3.1.1 Polycaprolactone (PCL) melt electrowriting

Fibrous scaffolds with a fibre diameter of 10 μm, square apparent pore size of 50 μm, and layer offset of 300⍰μm were fabricated on a custom built MEW apparatus as previously described [28]. We have demonstrated that this fibrous scaffold architecture is optimal for MSC osteogenesis [28]. Briefly, heated air at a temperature of 90°C was circulated around a syringe to melt polycaprolactone (PCL) (Sigma Aldrich 440744, average Mn 80,000), with air pressure being used to extrude the polymer through a 21G needle with high voltage applied at a distance of 15 mm from a grounded aluminium collector plate. Fibres were deposited with a 90° offset between subsequent layers to result in a square pore shape. Scaffolds for control (C) groups were used without further mineral modification.

#### 3.1.2 Plate hydroxyapatite (pHA) coating

This coating was carried out via a saturated simulated body fluid (SBF) solution, as described by Martine *et al* [48]. Reactions were carried out in 50ml conical tubes, with reagent volumes being maintained at 40 ml at each step of the process and 6 scaffolds of dimension 3 x 3 cm being processed per tube. Scaffolds were immersed in 70% ethanol for 15 min under vacuum and treated with 2M NaOH for 45 min at 37°C following a 5 min vacuum treatment to remove air bubbles. Scaffolds were rinsed 5 times in MilliQ water and immersed in 10x SBF, previously brought to a pH of 6 for 30 min at 37°C following a 5 min vacuum treatment. The coating step in 10x SBF was repeated a further two times minus the vacuum treatment. Scaffolds were then treated with 0.5M NaOH for 30 min at 37°C, rinsed 5 times in MilliQ water and allowed to dry overnight. Scaffolds treated with this coating method are defined as plate hydroxyapatite (pHA).

#### 3.1.3 Nano-needle hydroxyapatite (nnHA) coating

We developed a nano-hydroxyapatite coating to coat scaffolds with nano-needle mineral units. Sample pre- and post-processing was carried out as per the calcium phosphate coating procedure, with the 10x SBF treatment step being substituted for alternative calcium and phosphate solutions [49, 50]. The calcium solution was made with 0.05M calcium chloride dihydrate (Sigma Aldrich C7902) in MilliQ water. The phosphate solution was made with 0.03M sodium phosphate tribasic dodecahydrate (Sigma Aldrich S7778) and 0.01M sodium hydroxide in MilliQ water. The entire coating procedure is outlined as below. Scaffolds were immersed in 70% ethanol for 15 min under vacuum and treated with 2M NaOH for 45 min at 37°C following a 5 min vacuum treatment to remove air bubbles. Scaffolds were rinsed 5 times in MilliQ water and added to 20 ml of the calcium solution. 20 ml of the phosphate solution was slowly added to the calcium solution, and scaffolds were incubated for 30 min at 37°C following a 5 min vacuum treatment. This coating procedure was repeated a further two times minus the vacuum treatment. Scaffolds were then treated with 0.5M NaOH for 30 min at 37°C, rinsed 5 times in MilliQ water and allowed to dry overnight. Scaffolds treated with this coating method are defined as nano-needle hydroxyapatite (nnHA).

#### 3.1.4 Hydroxyapatite incorporated (iHA) PCL melt electrowriting

A nHA-PCL blend was created by incorporating a 3.5wt% of nano-hydroxyapatite (Sigma Aldrich 677418) into the polymer. PCL and nano-hydroxyapatite with a total mass of 1 g was added to a glass vial, with 10 ml of chloroform being added under stirring until dissolved. Particles were dispersed in the solution by sonication for 1 h, after which it was poured into a glass dish and left overnight for chloroform to evaporate. The resulting film was formed into pellets which were loaded into 1 ml BD-Braun syringes. The syringes were heated for 1 h at 110°C, and centrifuged at 4,000g for 60 s to remove air. The previous heating and centrifugation steps were repeated for a total of 8 cycles, or until a homogenous blend was present in the syringe with no air bubbles. MEW processing parameters were optimised to facilitate controlled deposition of PCL-nHA blends, with temperature increased to 110°C and voltage increased by approximately 0.5 kV. Scaffolds for this modification method are defined as fibre incorporated hydroxyapatite (iHA).

### 3.2 Scaffold characterisation

#### 3.2.1 SEM imaging and Energy-dispersive X-ray spectroscopy

For SEM imaging, samples were prepared for imaging by coating with gold/palladium for 40 s at a current of 40 mA. For analysis with energy-dispersive X-ray spectroscopy (EDX), scaffolds were carbon coated and analysed at a voltage of 15kV in a Zeiss ULTRA plus SEM with an 80mm^2^ Oxford Inca EDX detector. To investigate approximate calcium/phosphorus atomic ratios for each group, spectra were acquired on scaffolds for 30 s (n=6, 2 technical replicates per scaffold). Element maps were constructed by taking 2 frames at a resolution of 256 x 192 with map dwell of 4000 μs and linescan dwell of 2000 μs.

#### 3.2.2 X-ray diffraction

To obtain powder samples for x-ray diffraction (XRD) analysis, 10 x SBF was allowed to precipitate in solution at a pH of 6, while for the nnHA group, calcium and phosphate solutions were mixed together and allowed to precipitate. Solutions were centrifuged at 5,000g for 40 min to collect the precipitate, which was then placed in dishes and dried for 4h at 50°C in an oven. The resulting mineral was ground into a powder using a mortar and pestle. The hydroxyapatite powder used for iHA was used for XRD analysis without any further processing. Samples were loaded in a Brucker D8 ADVANCE powder diffractometer and run for 1 h from 5 - 80° in the 2θ range with a monochromatic Cu-Kα radiation source. Values for crystallinity and crystallite size were calculated using DIFFRAC.SUITE EVA 4.1.1. Crystallite size was calculated from taking 5 measurements of the (211) peak in pHA and iHA groups. In the iHA group, crystallite size was calculated from the peak in the position of (211), (112) and (300), due to the broad peaks resulting in their merging.

#### 3.2.3 Calcium staining

Samples were taken from scaffolds using a 2mm biopsy punch (n=5) and incubated with 1% alizarin red s (Sigma Aldrich, A5533) in distilled water for 10 min at a pH of between 4.1 - 4.3. Samples were rinsed 3 times with water and allowed to dry prior to imaging. To quantify bound stain, 400 μl of 10% acetic acid was applied and samples incubated at room temperature for 30 min while shaking at 150rpm. The acetic acid was added to 1.5 ml tubes, vortexed vigorously and heated to 85°C for 10 min. Tubes were transferred to ice for 5 min, centrifuged at 20,000g for 15 min, and 300 μl of the supernatant was added to new tubes along with 120 μl of 10% ammonium hydroxide. Standards were made with dilutions of alizarin red solution in distilled water, with the pH of each adjusted to between 4.1 - 4.3. Samples and standards were read at 405 nm in a 96-well plate.

#### 3.2.4 Water contact angle

The water contact angle of all scaffold groups was quantified using a FTA125 contact angle analyzer (First Ten Angstroms Inc.). Samples were prepared using an 8mm biopsy punch (n=6-7). All measurements were taken 10 frames after contact of the water with the sample.

#### 3.2.5 Tensile testing

Rectangular samples (n=4) of dimension 5 x 20 mm were taken from scaffolds and used for tensile testing in a Zwick Z005 with 50⍰N load cell (A.S.T. - Angewandte System Technik GmbH). Samples were pre-loaded to 0.002⍰N at a speed of 1⍰mm/min, and then loaded at a speed of 10⍰mm/min with a time save interval of 0.1⍰s. Load-displacement curves were graphed and used to determine the stiffness, toe region stiffness and yield force for each group.

### 3.3 Human bone marrow mesenchymal stem/stromal (hMSC) cell culture

Scaffolds were punched to a diameter of 8 mm and UV sterilised for 20 min on each side before being mounted in stainless steel holders and placed in 24-well plates. Scaffolds were then pre-wet in a graded ethanol series of 100%, 90% and 70% for 20 min each before being washed three times in phosphate buffered saline (PBS). They were then incubated overnight in DMEM with 10% FBS at 37°C. Human bone marrow mesenchymal stem/stromal cells (hMSCs) were isolated from bone marrow (Lonza, US), trilineage potential verified (data not shown), and seeded at a number of 10,000 cells per scaffold. Scaffolds were transferred to new well plates after 24 h, and cultured in osteogenic medium (OM) from day 3, which consisted of 10% FBS DMEM supplemented with 100⍰nM dexamethasone, 10⍰mM β-glycerol phosphate and 50⍰μg/ml ascorbic acid. Medium was changed every 3.5 days.

### 3.4 Proliferation

At days 1,7,14 and 21, scaffolds were added to 100⍰μl lysis buffer (n=4) containing 0.2% Triton X-100, 1⍰mM Tris pH8, with phenylmethylsulfonyl fluoride (PMSF) being added at a ratio of 1:200 just before use, and stored at −80°C. Before DNA quantification, samples underwent sonication for 60 s and subjected to three freeze-thaw cycles in liquid nitrogen before being stored on ice. DNA content was quantified using a Quant-iT™ PicoGreen™ dsDNA Assay Kit (Invitrogen, P7589), with excitation and emission wavelengths of 485⍰nm and 528⍰nm respectively. The DNA content in 10,000 hSSCs seeded and pelleted in centrifuge tubes was also quantified to calculate the total number of cells present on scaffolds.

### 3.5 Characterisation of hMSC osteogenic differentiation

#### 3.5.1 Intracellular alkaline phosphatase (ALP)

Intracellular ALP was quantified at days 14 and 21 (n=4). 50 μl of 5mM pNPP was added to wells along with 10 μl of cell lysate and 70μl MilliQ water in a 96-well plate. Standards were comprised of serial dilutions of p-Nitrophenyl phosphate (pNPP, Sigma Aldrich, N1891) with 10μl of 43μM ALP enzyme (Sigma Aldrich, P6774) being added to each. Plates were incubated for 1 h in the dark at room temperature, and reactions were then stopped using 20μl of 3M NaOH. Absorbance was read on plates at 405 nm, and ALP activity was calculated as the amount of pNPP generated as a function of sample volume and reaction time.

#### 3.5.2 Collagen production

Scaffolds were cultured up to 21 days before evaluating collagen content. Cell-scaffold constructs were rinsed in PBS, fixed in 10% neutral buffered formalin for 15 min and rinsed again in PBS before storage at −20°C. Scaffolds were stained with 200μl of 1mg/ml of Direct Red 80 (Sigma Aldrich, 365548) in a saturated aqueous picric acid solution for 1 h with shaking at 150 rpm. Scaffolds were then washed twice with 0.5% acetic acid and allowed to dry overnight before imaging. To quantify collagen content, 500μl 0.5M NaOH was added to wells under shaking until stain was dissolved, and solutions were added to 1.5 ml tubes. Tubes were centrifuged at 14,000g for 10 min to pellet debris. Standards were made by adding direct red staining solution to 8μl of collagen I (Corning, #354249) before centrifuging at 14,000g for 10 min and re-suspending the collagen in 500μl 0.5M NaOH. The absorbance of samples and standards were read at 490 nm in 96-well plates.

#### 3.5.3 Calcium production

Cell-scaffold constructs were investigated for total calcium content after 21 days using 1% alizarin red S solution as previously described for calcium staining in scaffold characterisation. Cell-free scaffolds were also cultured up to 21 days to determine the contribution of total calcium from cell mineralisation versus the calcium present due to mineral modification of scaffolds.

### 3.6 BMP2 functionalisation of MEW scaffolds

#### 3.6.1 BMP2 adsorption onto scaffolds

Scaffolds were prepared, sterilised and pre-wet in a graded ethanol series as previously described in section 3.3 before being washed three times in PBS. Recombinant human BMP2 (Peprotech 120-02) was diluted to 50 μg/ml in PBS, with 20 μl of solution containing a total of 1 μg BMP2 being placed on each scaffold and incubated for 4 h at room temperature. BMP2 was then removed and scaffolds were allowed to dry overnight.

#### 3.6.2 BMP2 release kinetics

Scaffolds were placed in 48-well plates (n=4) and 200 μl of PBS added to each. At each time-point, PBS was removed and 1% BSA added to result in a final BSA concentration of 0.1%. 200 μl of fresh PBS was then added to each well. Samples were stored at −80°C until BMP2 quantity was determined via ELISA (R&D Systems DY355-05). Cumulative BMP2 release was studied at time-points 1 h, 6 h, and days 1, 3, 7, 14 and 21.

#### 3.6.3 Characterisation of osteogenic differentiation

After BMP2 adsorption, 10,000 hMSCs were seeded onto scaffolds (n=4), and cultured up to 21 days after which DNA content was investigated to determine total cell number. Scaffolds were also cultured up to 21 days to assess osteogenic potential in terms of ALP activity, collagen production and calcium production as described previously.

### 3.7 Statistical analysis

Tensile testing and material characterisation data is presented in terms of average and standard deviation. Subsequent biological data is presented in terms of average and standard error of the mean. Statistical analysis was performed using one-way ANOVA and Tukey’s multiple comparison post-test. Data from the BMP2 loading study was analysed using two-way ANOVA and Bonferroni’s multiple comparison test.

## 4 Results

### 4.1 MEW fibre topography is significantly altered by bulk and surface modification

The topography of 90° orientate fibrous MEW scaffolds was investigated following modification with calcium phosphate based materials, where significant changes were identified with minimal influence on macroscopic scaffold morphology. Control fibres have a diameter of 9.1 μm (SD = 1.0), with relatively featureless surface topography (Figure 1A). The pHA coating protocol yields a rosette-like arrangement of plates of average diameter 555.3 nm (SD = 193.2) and thickness 26.0 nm (SD = 8.6) (Figure 1B). Total fibre diameter including the coating is 13.5 μm (SD = 2.6), giving an average coating thickness in the pHA group of 2.2 μm. In stark contrast to this, the nnHA coating protocol results in the formation of nano-needles on the fibre surface (Figure 1C), with a length and diameter of 100.0 nm (SD = 29.0) and 37.0 nm (SD = 7.1) respectively. These needles form occasional aggregates which range in diameter from 150 - 500 nm. Total fibre diameter shows a minimal increase compared to the control with a value of to 10.2 μm (SD = 1.0), yielding a total coating thickness in this group of 0.55 μm. The incorporation of HA within fibres in iHA during the MEW process yields minimal changes in surface topography, with the majority of visible particles being seen as bulges just below the fibre surface, while several can also be seen to protrude above the surface with no covering of PCL (Figure 1D). Particles have an average diameter of 147.2 nm (SD = 78.5), as measured within PCL fibres. The incorporation of HA also has an influence on the quality of the MEW process, with periodic pulsing of the Taylor cone resulting in a larger fibre diameter distribution (fibre diameter = 11.8 μm, SD = 4.2).

**Figure 1.**
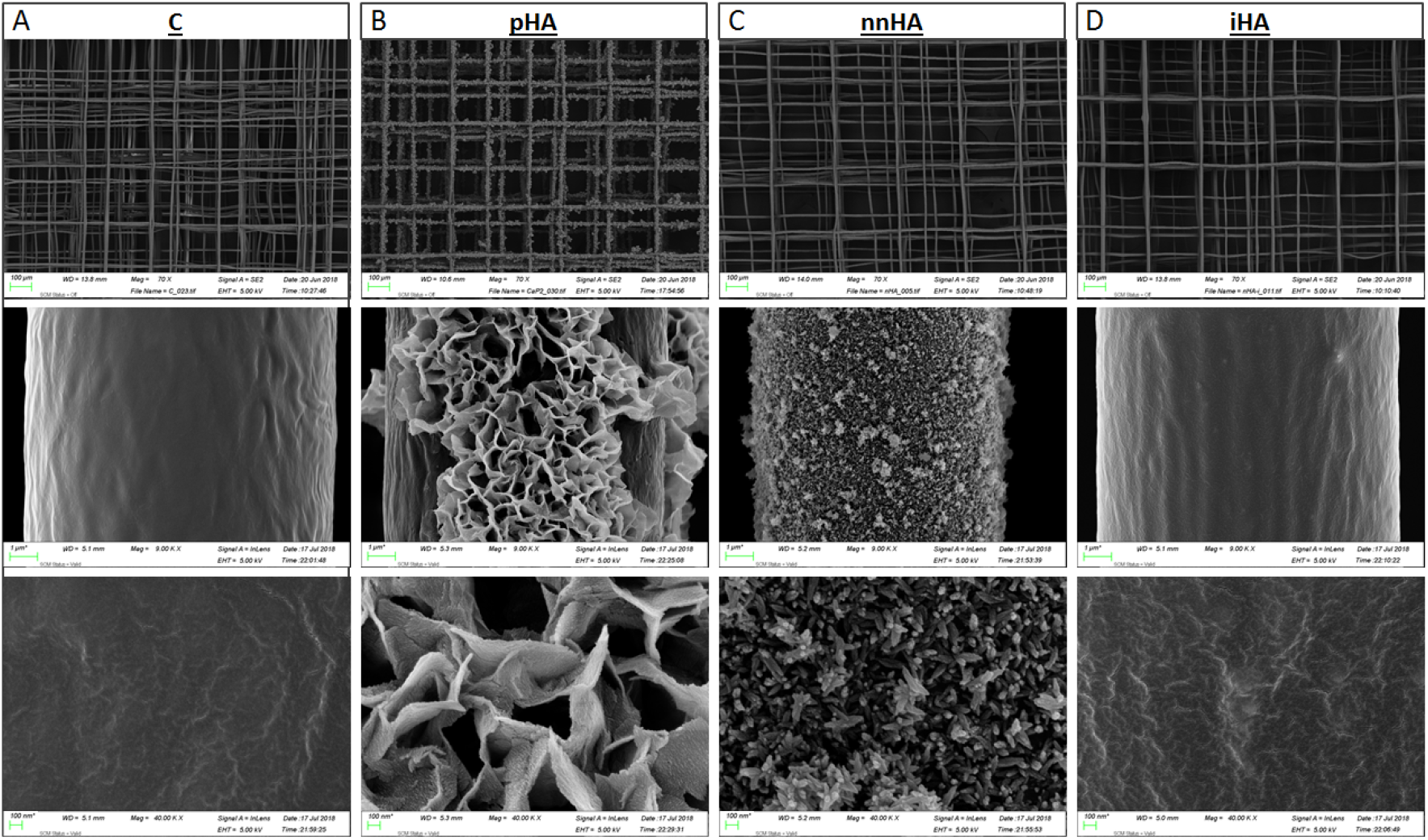
SEM imaging of scaffold groups illustrating the "plate" coating morphology in pHA, "needle" morphology in nnHA, and the presence of HA nanoparticles both within and protruding above the surface of the fibres in iHA.

### 4.2 CaP treatments modify surface chemistry

#### 4.2.1 Element analysis

The coating chemistry and distribution in fibres was investigated via EDX analysis. The presence and distribution of calcium and phosphorous was first investigated via element mapping where, as expected, there was no detection in the control (Figure 2A). A consistent distribution of both elements was seen in pHA and nnHA coated fibres, while a sparse distribution in the iHA group confirmed the presence of HA particles within fibres. Carbon and oxygen were present in all groups from the PCL, as well as calcium and phosphorous as seen in the spectra of all groups except the control (Figure 2C, merge Figure 2D). A key difference in the pHA group was seen with the presence of sodium and magnesium in this coating. Ca/P ratios were investigated (Figure 2D) and were found to be close to stoichiometric hydroxyapatite with a ratio of 1.66 for the pHA group. The iHA group was investigated using both MEW scaffolds with HA incorporated, as well as the stock powder, with both having a resulting ratio of 1.76. The nnHA coating group deviated slightly from these findings, with a more calcium-rich HA with a ratio of 1.97 being identified.

**Figure 2.**
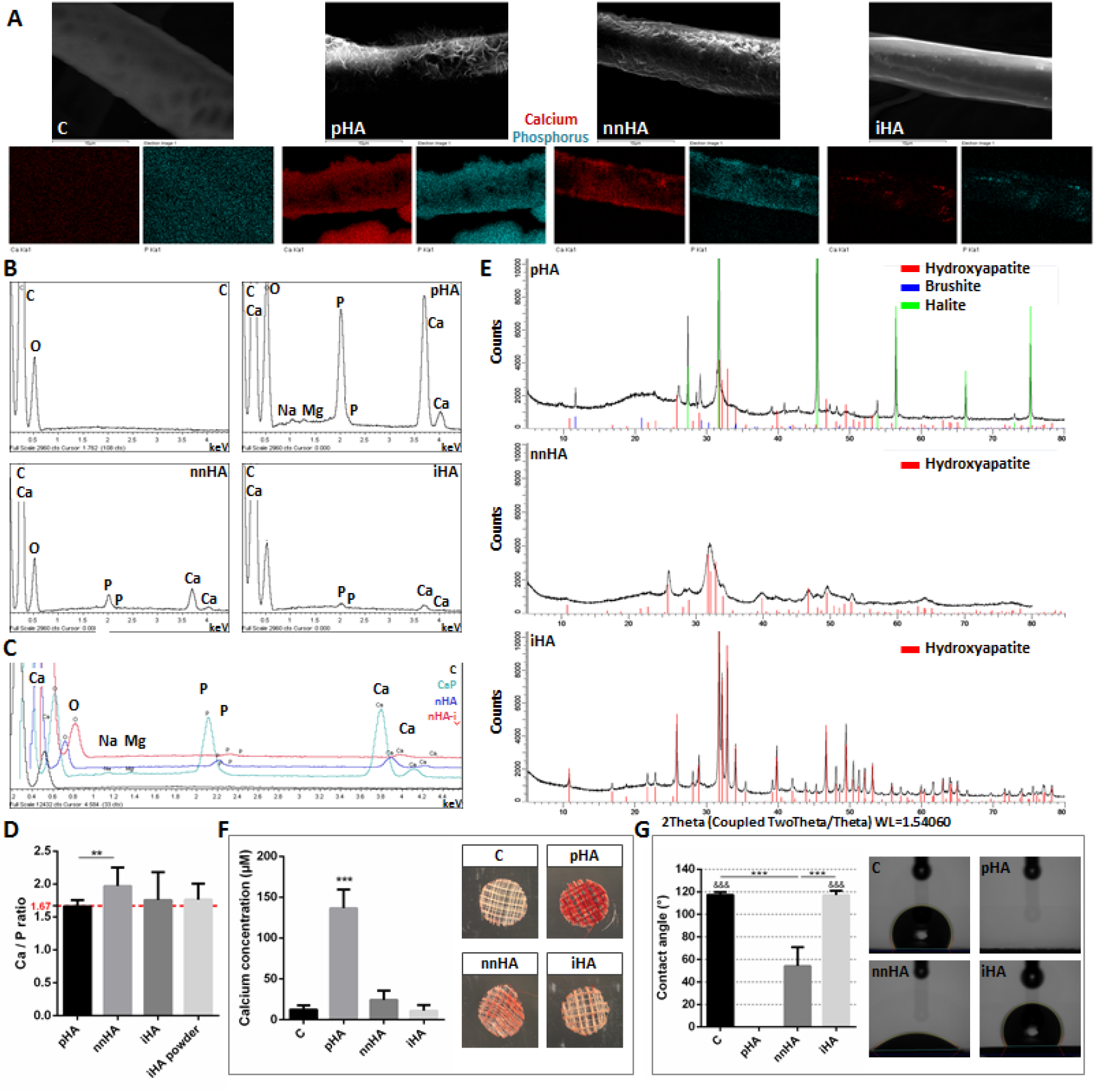
Chemical and physical characterisation of MEW-hydroxyapatite scaffolds. Element mapping of SEM images via EDX analysis illustrating the distribution of calcium and phosphorus in each group (A). Representative EDX spectra of all groups (B) and merge of spectra (C). Calcium to phosphorus ratios of EDX spectra (n=6 scaffolds, n=2 technical replicates) (D). XRD analysis of powder representative of CP, nnHA and iHA groups (E). Alizarin red staining of scaffolds (n=5) (F). Water contact angle illustrating increased hydrophilicity after hydroxyapatite coating (n=6-7) (G).

#### 4.2.2 Crystal structure of surface and bulk modifications

XRD analysis was further conducted to characterise the crystal structure between groups (Figure 2E). Brushite, hydroxylapatite and halite were identified in the pHA group, in close agreement with previous literature [51], with a crystallinity of 61.3%. The nnHA and iHA groups were both shown to be composed solely of hydroxyapatite, with crystallinity of 52.5% and 69.3% respectively. The broader peaks in the nnHA group are indicative of a smaller crystallite size, which was found to be 8.5 nm. This is in comparison to the larger crystallite sizes in pHA and iHA, which were measured at 105.8 nm and 75.0 nm respectively.

#### 4.2.3 Calcium content of PCL modifications

The total calcium content as determined via quantification of alizarin red staining was investigated following scaffold modification with calcium phosphate minerals (Figure 2F). The calcium detected in C is minimal, and is due to slight background staining of the PCL. The greatest amount of calcium was found on pHA where nearly 150 μm was detected and was significantly greater than all other approaches. nnHA contained the second largest quantities of calcium but was still 6-fold less than that with pHA and was not significantly different to iHA. The calcium content detected in iHA is only marginally greater than that detected from background staining (C) due to the majority of HA particles being distributed throughout the bulk of the fibres.

#### 4.2.4 Surface treatment enhances material hydrophilicity

The hydrophilicity of groups was investigated and found to correlate closely with their calcium content (Figure 2G). This is seen in the complete spreading of water on pHA, which has the greatest calcium content. The nnHA group also exhibited a significant degree of hydrophilicity, with a mean contact angle of 56.0°. The C and iHA groups were both shown to be hydrophobic in comparison, with contact angles of 117.3° and 117.1° respectively.

4.3 Mechanical properties are enhanced via HA-incorporation into PCL MEW fibres

Incorporation of HA into PCL in iHA is seen to increase its stiffness and yield force, while all coating protocols have minimal influence on mechanical behaviour (Figure 3A,B). Stiffness in the linear elastic region between 0.1 - 0.3 N is greatest in iHA scaffolds (Figure 3C), while stiffness in the toe region is not seen to be significantly altered between groups (Figure 3D). Yield force of iHA scaffolds is also significantly enhanced compared to all other groups, with an average fold increase of 1.4.

**Figure 3.**
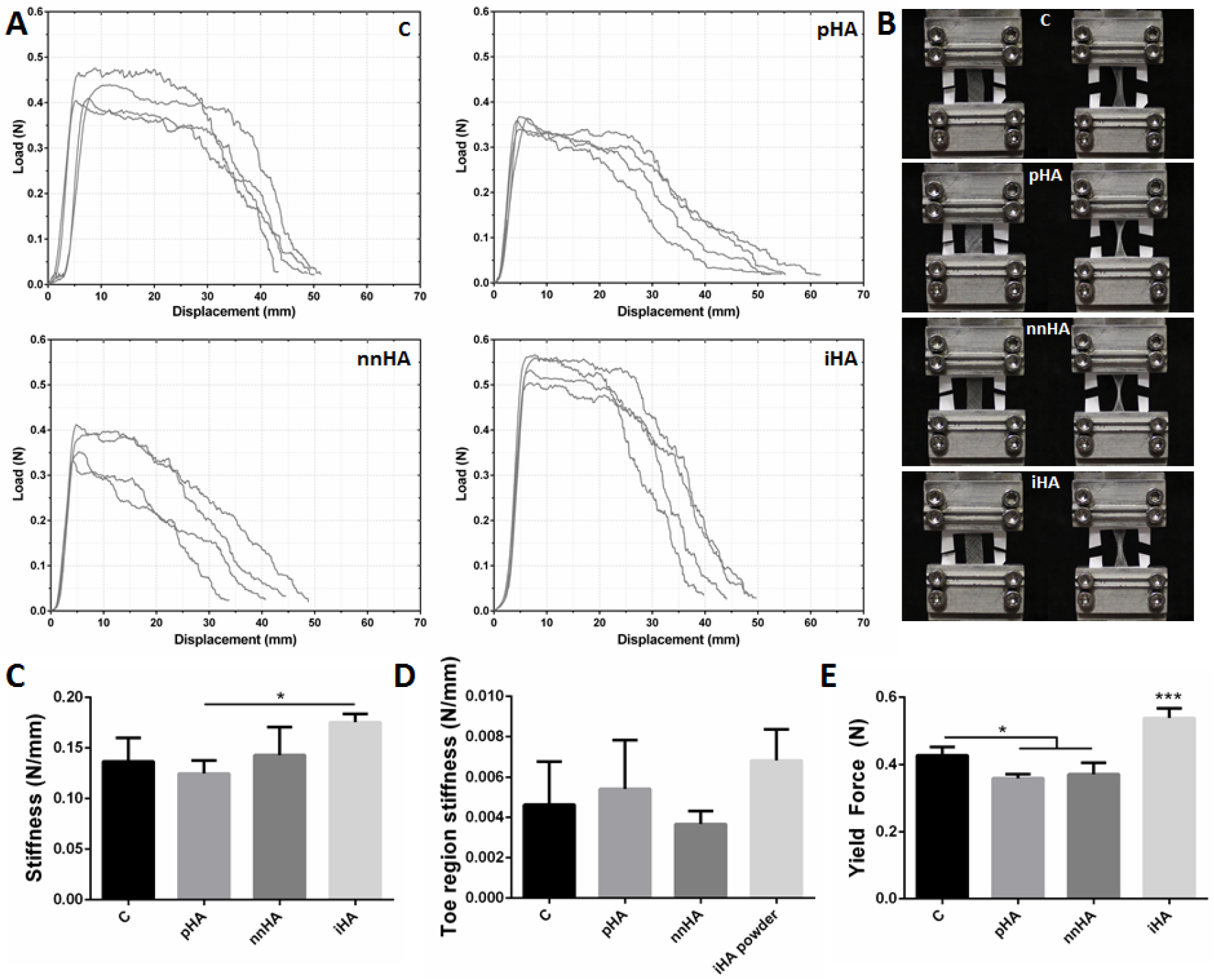
Mechanical characterisation of scaffolds via tensile testing. The incorporation of HA into fibres (iHA) can be seen to alter the mechanical properties of PCL micro-fibres (A). Scaffolds before tensile testing, and after being loaded to approximately 50% of the yield strain (B). Significant increases in stiffness seen in iHA (C), with no change in toe region stiffness (D). Yield force of iHA scaffolds is also significantly increased (E). (n=4).

### 4.4 Attachment and proliferation of hMSCs

Cells adhere to scaffolds with an average seeding efficiency of 25%, with approximately 1,100 - 3,400 cells remaining as quantified after 24 h post seeding. Adherent cells were found to branch across fibres resulting in elongated. Adherent hMSCs were commonly found branching across adjacent fibres as has been previously demonstrated with this fibre architecture [28]. Interestingly the pHA coating resulted in a slightly altered cellular morphology will hMSCs adhering and spreading across the deposited mineral, masking the underlying fibre architecture (Figure 4A). The initial cell seeding efficiency is lowest in the coated scaffold groups; pHA and nnHA. This trend is seen to level off by D7 however, where cell number reaches the initial seeding quantity at a number of 10,000 per scaffold. Cells then proliferate 2-fold per week for the remainder of the study, with an average of 24,000 cells at D14 and 41,000 cells at D21. While there are no significant differences between groups at any time-point, a trend of reduced cell number in the pHA scaffold group is seen at D14 and D21.

**Figure 4.**
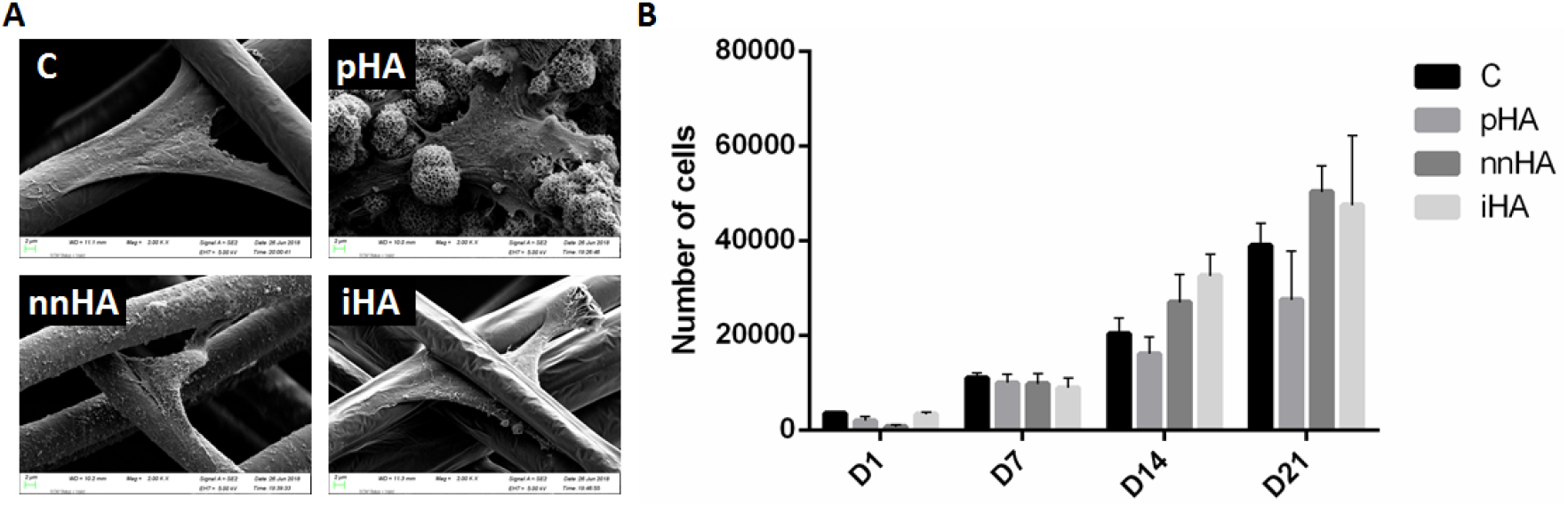
SEM imaging of cell seeded scaffolds after 24 h culture (A) Cell proliferation as quantified by DNA content. Proliferation in pHA scaffolds can be seen to be attenuated D14 compared to other scaffold groups (B). (n=4).

### 4.5 Surface treatment significantly enhances hMSC osteogenesis

#### 4.5.1 ALP activity

No significant trends were identified in hMSC ALP expression between each scaffold at D14 and D21. Total ALP activity can however be seen to be greater in nnHA and iHA groups at these time-points (Figure 5A). Of note is the 2.2-fold increase in ALP activity in nnHA at D21 compared to D14, in contrast with all other groups which remain constant between these time-points. Normalising these results to cell number reveals no further trends in ALP expression (Figure 5B), which remains relatively constant between groups.

**Figure 5.**
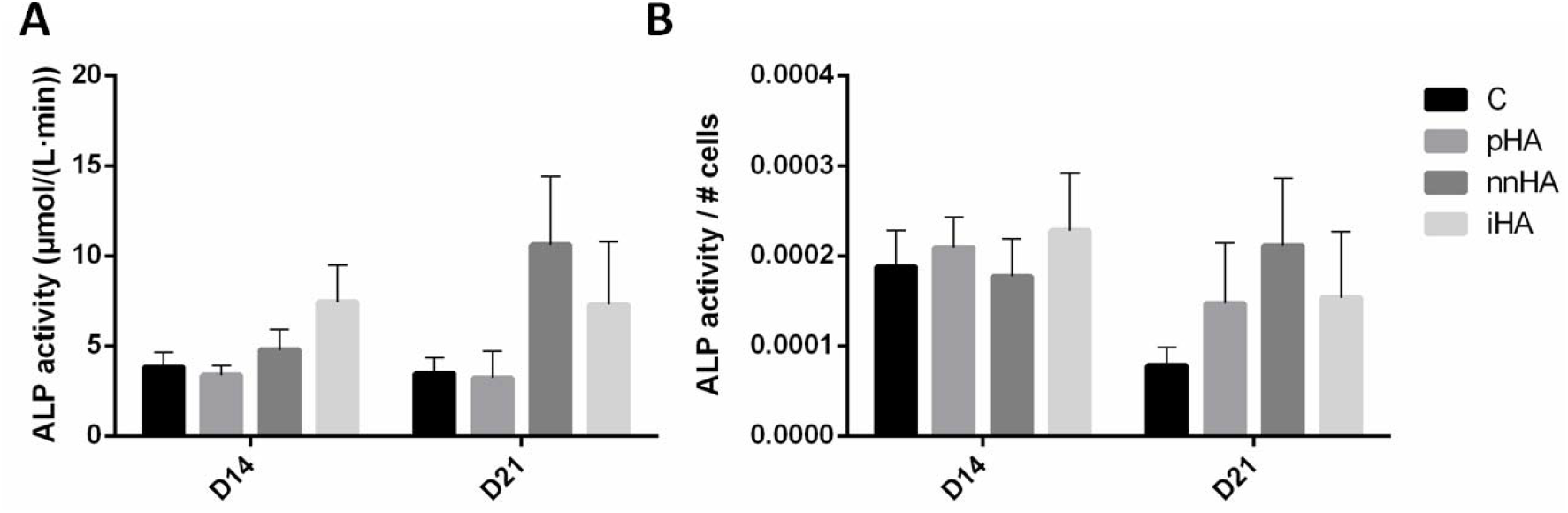
ALP activity in hMSCs at D14 and D21. No significant changes in total ALP activity (A) or ALP activity normalised to cell number (B) were identified. (n=3-4).

#### 4.5.2 hMSC Collagen production

Collagen production is shown to be enhanced in coated scaffold groups (Figure 6). Total collagen quantity is greatest in pHA and nnHA at *D21*, with a 1.7 and 1.5-fold increase respectively compared to control scaffolds. This effect is further magnified in the pHA group when normalising results to cell number (Figure 6C), which is significantly greater than other groups. Collagen content per cell is 3.4-fold greater in pHA than control, and over 2.5-fold greater than nnHA and iHA groups.

**Figure 6.**
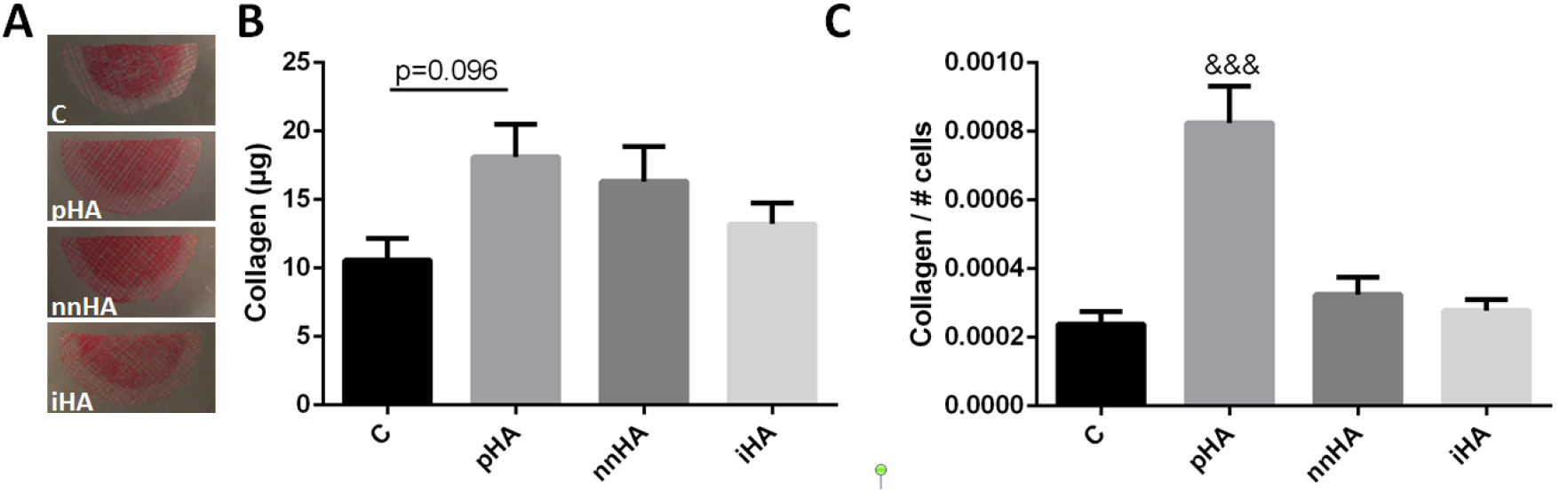
Collagen production at D21 (A). Total collagen is enhanced in the pHA group compared to C (A), while a significant increase compared to all groups is seen when collagen is normalised to cell number (B). (n=4). & = statistical significance compared to all other groups (&&& = p<0.001).

#### 4.5.3 hMSC Calcium production

As with previous outputs of osteogenic differentiation, mineral production is seen to be enhanced in the coated scaffold groups (Figure 7A). These results encompass both mineralisation due to calcium production by cells, as well as the calcium present due to the treatments prior to cell seeding. Total mineral in pHA was 6.7-fold greater than C and iHA (Figure 7B). The greatest concentration following 21 days culture was found on scaffold nnHA, with this being over 12-fold greater than the calcium content in C and iHA and 1.8-fold greater than pHA. These findings are further enhanced when considering calcium due to cell mineralisation alone. The cell contribution to mineral in pHA was 59%, with the remainder the mineral from the scaffold coating. In stark contrast, 92% of the mineral in nnHA is contributed by the cells. Considering this cell mineralisation alone, nnHA is over 14-fold greater than C and 2.9-fold greater than pHA. When total calcium is normalised to cell number at D21 (Figure 6C), calcium in pHA and nnHA groups are seen to be significantly greater than others, with an over 10-fold increase in mineral production per cell in pHA and over 12-fold increase in nnHA when compared to untreated control PCL scaffolds.

**Figure 7.**
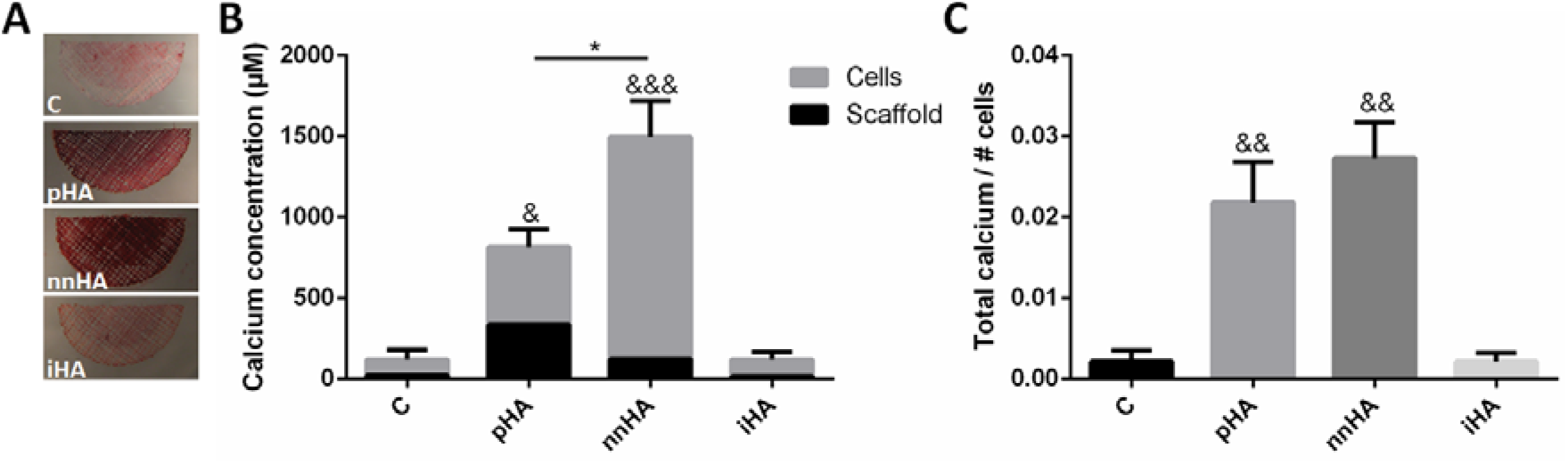
Calcium content at D21 with contributions from scaffold and cell mineralisation (A). Total calcium content is significantly enhanced in nnHA and pHA groups (B). Calcium content normalsied to cell number is also enhanced in these groups compared to C and iHA (C). (n=4). & = statistical significance compared to C and iHA groups (& = p<0.05, && = p<0.01, &&&= p<0.001). * = p<0.05.

### 4.6 Coating dissolution after long term culture

After 21 days in culture, the pHA coating was found to be largely intact as seen via SEM imaging, with minor changes seen in a more rough surface topography (Figure 8B) compared to at D1 (Figure 8A). In stark contrast to this, the nnHA coating is seen to have undergone considerable dissolution, with loss of the nano-needle topography and formation of a coating of nano-spheres with diameter of approximately 5-20 nm. Consistent fracturing of this coating can also be seen. The iHA group is also considerably altered, with extensive precipitates seen at the fibre surface in contrast to the relatively smooth fibre topography at D1.

**Figure 8.**
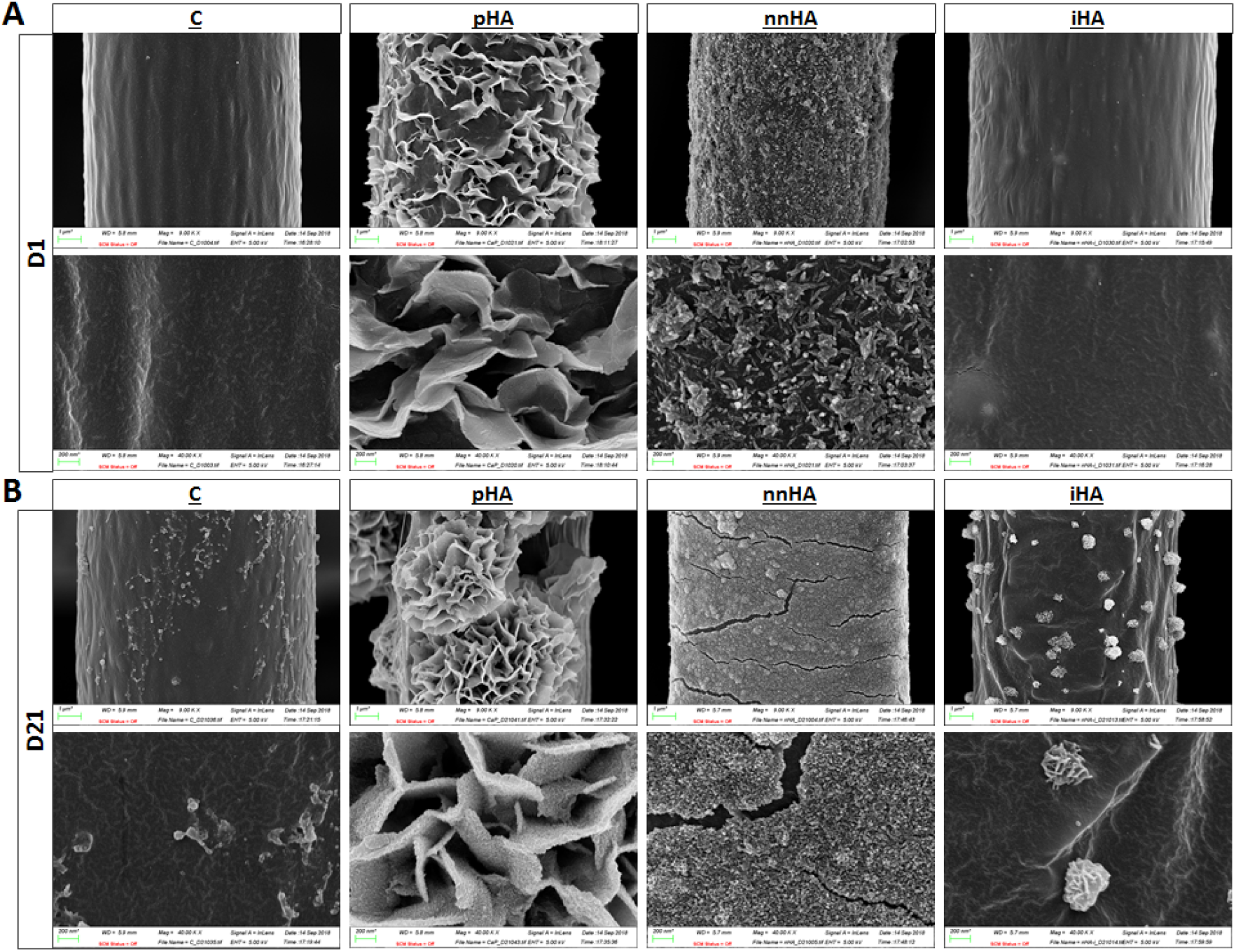
Fibre appearance after day 1 (A) and day 21 (B) in culture. Plates in the pHA group appear more rough at D21, with morphology otherwise unchanged. The nnHA coating is significantly altered, with the initial nano-needles becoming spherical and smaller in nature, and the coating exhibiting widespread fracturing. A greater amount of mineral precipitation on the fibre surface is seen in iHA at D21, where the near-surface HA appears to act as sites for mineral nucleation.

### 4.7 Mineral architecture mediates BMP2 controlled release and further enhances hMSC osteogenesis

In order to investigate the capacity of mineral architectures to bind, stabilise, and release growth factors, scaffolds were loaded with BMP2 and release kinetics and stem/stromal cell osteogenesis investigated. There is an initial burst release of BMP2 from the C, pHA and iHA materials, with a more sustained initial controlled release from nnHA (Figure 9A). The majority of the BMP2 is released by C after 7 days with a total amount of approximately 50%, where the curve is seen to plateau with minimal further release up to day 21, indicating a lower level of adsorption of the protein with reduced affinity to the scaffold (Figure 9B). At 7 days, release percent from pHA and iHA is 65% and 61% respectively, in contrast to nnHA, with a value of 46%. Between D7 and D21, nnHA continues to maintain the greatest level of controlled release at a rate of 9.8% per week.

**Figure 9.**
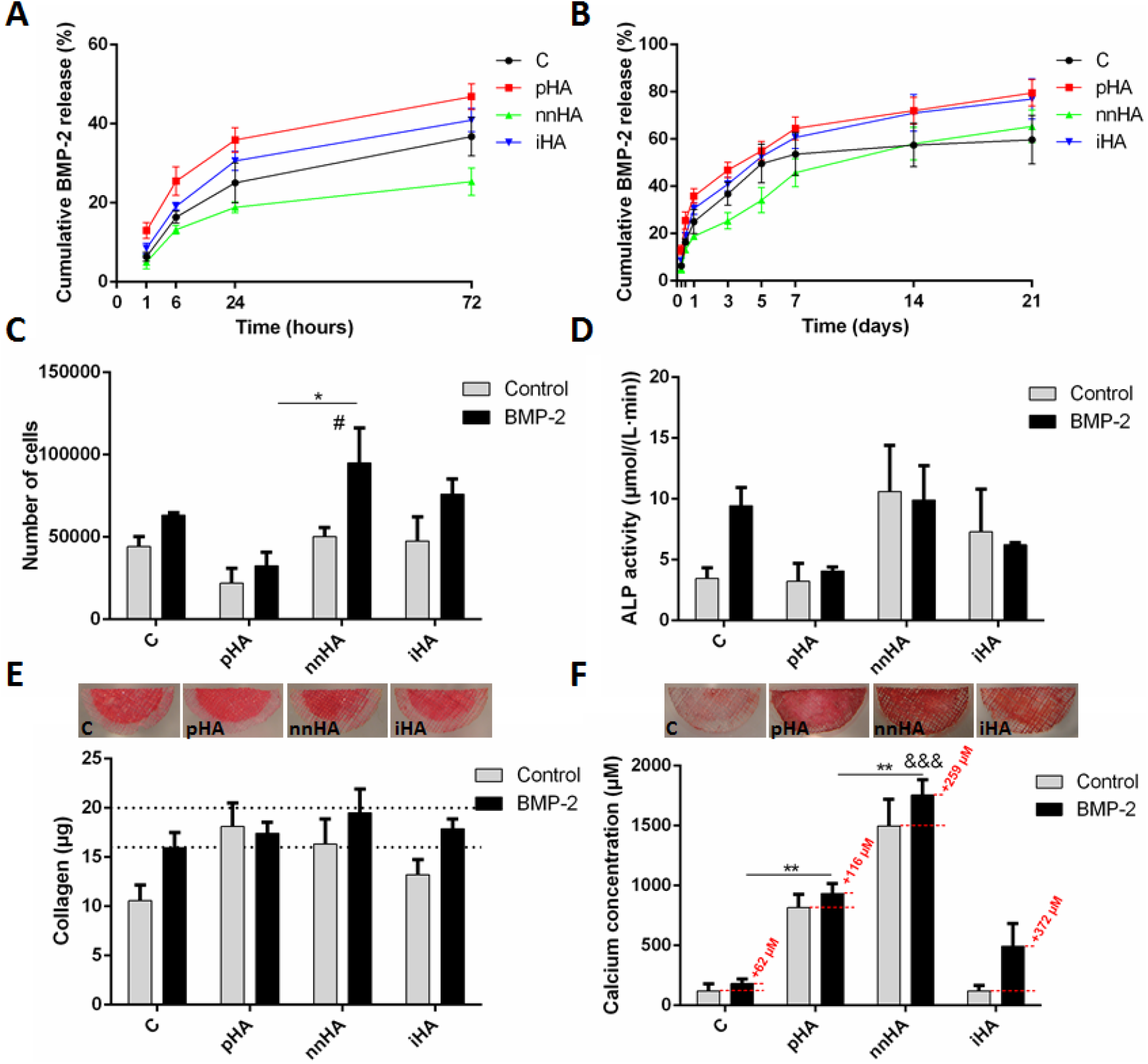
Cumulative BMP2 release up to D3 (A) and up to D21(B) (n=4). Number of cells (C), ALP activity (D), collagen content (E) and calcium content (F) with BMP2 treatment at D21 compared to data from untreated scaffolds as presented in previous figures. * = statistical significance via one-way ANOVA and Tukey multiple comparison test within BMP2 group only. & = statistical significance vs C and iHA groups with same method as previously. # = statistical significance between Control and BMP2 in the nnHA group via two-way ANOVA and Bonferroni’s multiple comparison test.

The controlled release of BMP2 from the MEW scaffolds is seen to enhance cell proliferation in each group. Of particular note is the significant 1.9-fold increase in cell number on the nnHA scaffolds at day 21 compared to control nnHA scaffolds, in addition to significantly enhanced DNA in BMP2 treated nnHA vs pHA, with a fold change of 2.9 (Figure 9C). The effect of BMP2 delivery on the osteogenic differentiation of hMSCs was investigated via ALP activity, collagen and mineral deposition as before. ALP increases were only seen in the C scaffolds, however as with all other groups, this was not significant (Figure 9D). Collagen content was also investigated, with the greatest increases in C and iHA compared to control scaffolds (Figure 9E). Overall collagen content was between 16-20 μg in all groups, with greatest collagen content in nnHA after 21 days. Finally, mineralisation is further enhanced in CP BMP2 functionalised scaffolds, with the greatest changes seen in the nnHA and iHA groups, with increases of 259 μM and 372 μM respectively (Figure 9F). In summary, the nnHA group exhibits the greatest potential for modification with additional factors, as seen by the greatest level of sustained protein release, significantly enhanced DNA content, and the greatest levels of collagen content and mineralisation content following 21 days in culture.

## 5 Discussion

Due to a lack of bone autograft supply, bioinspired synthetic alternative materials are required to enhance bone regeneration in critically sized defects. Cell behaviour and function are mediated by microenvironmental biophysical and biochemical cues, with the architecture and chemistry of the surrounding extracellular matrix being fundamental in this regard. Bone at a cellular level consists of arranged fibres with a cross-fibrillar mineral phase made up of curved nano-sized needle shaped crystals. Recapitulating this unique multiscale architecture may represent an innovative approach to develop effective bioinspired materials to promote bone regeneration. In this study, we utilise a 90° fibrous MEW scaffold architecture, which we have previously optimised for human MSC osteogenesis [28], and built on this foundation by developing biomimetic mineral functionalisation strategies consisting of either intrafibrillar or cross fibrillar configurations. Interestingly using a novel coating approach, we were able to grow nano-needle shaped HA (nnHA) crystals on the surface of the MEW fibres which closely mimic that found in bone. These bioinspired nnHA topographies significantly enhanced hMSC osteogenesis over traditional SBF plate HA (pHA) coatings or intrafibrillar HA (iHA) scaffolds. Moreover, given the protein stabilising properties of mineral, we demonstrated that nnHA facilitated a controlled release of BMP2 from the scaffold which further enhanced mineral deposition by stem/stromal cells. This study thus outlines a method for fabricating scaffolds with precise fibrous micro-architectures and bone mimetic nano-needle HA extrafibrillar coatings which significantly enhance MSC osteogenesis and therapeutic delivery and thus hold great promise for bone tissue regeneration.

The microarchitecture/topography, material chemistry, and mechanical properties of each MEW scaffolds is altered followed CaP modifications. The concentrated SBF pHA coating technique resulted in the formation of plates arranged in a "cauliflower" or "rosette" like coating morphology, consistent with previously published findings using this approach [51–53] and typical of the plate or "petal-like" morphology of apatite formed via supersaturated solutions [54]. In contrast, the nnHA coating procedure which we have developed resulted in a vastly different morphology consisting of a fine, extrafibrillar, nano-needle coating which form occasional aggregates, closely mimicking the fundamental mineral units of bone in terms of morphology and composition recently identified [4]. The incorporation of HA into PCL in the iHA group does not have a significant influence on fibre surface morphology, as seen by SEM and water contact angle measurements, however, mechanical properties are enhanced with this approach. EDX analysis was further conducted to investigate the elemental distribution and composition of modified fibres. The Ca/P ratios of all groups were within the range seen in native bone, which has typical Ca/P ratios varying between 1.34 - 2.17 with a median value of 1.69, as seen in samples from the femoral neck and the rib bones of healthy humans [55–57] (note: weight ratios in references were converted to atomic ratios). To further investigate the crystal structure of our scaffold groups, we performed an x-ray diffraction analysis. Interestingly, the broad peaks of the XRD spectra of nnHA closely mimic those of native bone. This is in comparison to the other groups which are more crystalline in nature. In summary, we have demonstrated that our nnHA coating procedure closely mimics the nano-architecture, composition and crystal structure of native bone.

Bone bioinspired architectures can significantly enhance MSC mediated bone formation, as we have demonstrated via long term in-vitro studies. Cell behaviour was investigated on modified scaffolds via proliferation and markers of osteogenic differentiation. Cell number was greatest in the nnHA group after 21 days, with the lowest number of cells in the pHA group. This is consistent with previous work demonstrating a trend of increased proliferation with reduced HA particle size [34]. We have also demonstrated a corresponding increase in ALP activity in nnHA, however, this result is not significant. It is known that osteoprogenitor cells initially produce ALP and secrete collagen type I, with further mineralisation occurring on this matrix at later time-points [58, 59]. We have shown previously that the fibrous architecture of scaffolds can have a significant influence on hMSC collagen production [28]. However, our current findings indicate that HA surface topography does not augment collagen production. Further investigation of calcium deposition demonstrated highly significant upregulation, particularly with nnHA coating. The majority of the contribution towards total calcium content is seen to be due to cell-based mineralisation, indicating the significant influence of nano-scale needles in this extrafibrillar coating in driving stem/stromal cell osteogenic differentiation. There are several likely factors for this result, in addition to the influence of the nano-particle size as discussed previously. Greater dissolution of this coating, as demonstrated by SEM imaging after 21 days in culture will yield a greater release of calcium and phosphate ions into the medium which can interact with cells and allow for reprecipitation, enhancing mineralisation [60]. In addition, an inverse relationship between crystallinity and mineralisation has also been shown [61], providing further evidence for the greater cell mineralisation on our nnHA group, which has the lowest crystallinity. This therefore suggests that the superior bone formation demonstrated with the nnHA coating is multifactorial, harnessing several mechanisms to maximise regeneration.

Nano-needle structured HA extrafibrillar coatings effectively bind, and slowly deliver stable BMP2, enhancing hMSC proliferation and bone formation. We investigated the protein binding and release capacity of these scaffold modifications via a BMP2 adsorption study. Recent studies have demonstrated that mineral can stabilise bound proteins, preventing conformational changes, maintaining biological activity for weeks during release and that this stabilisation is enhanced within nanostructured coatings [6]. The nnHA group was shown to facilitate the most sustained release of BMP2 over 21 days. Interestingly, this resulted in significantly enhanced proliferation in this group demonstrating biological activity, while markers of osteogenic differentiation were also marginally increased. A previous study has reported similar results, where continuous supplementation of media with BMP2 resulted in enhanced proliferation of hMSCs without initiating osteogenic differentiation [62]. However, many studies have also demonstrated the key role of BMP2 in stimulating osteogenesis [63–65]. We believe that due to the high degree of mineralisation indicating late stage differentiation, day 21 may be too late to capture elevated ALP expression, which is an early differentiation marker. Consistent with previous findings in this paper, we also believe that peak collagen levels for our scaffolds are being reached, as seen by the attenuated difference between groups with BMP2 treatment compared to untreated scaffolds. While no significant increases in calcium content were identified, it must be noted that the levels in nnHA are already 14fold greater than control scaffolds in the absence of BMP2 treatment, and the further increases in mineral with BMP2 treatment in nnHA and iHA groups represent a change several fold-times greater than basal calcium levels in untreated control scaffolds, which we have previously shown to represent significantly enhanced mineralisation compared to random scaffolds [28]. In summary, scaffolds with nnHA coating are shown to significantly enhance stem/stromal cell differentiation, with the controlled release of BMP2 loaded scaffolds further contributing to this effect.

## 6 Conclusions

In conclusion, we utilised MEW technology to create fibrous 3D micro-architectures and further modified these templates with a novel bioinspired extrafibrillar coating of needle shaped nano-sized crystals (nnHA). This bone mimetic fibrous and mineral architecture significantly enhanced human MSC osteogenesis over more established plate like mineral coatings. Moreover, extrafibrillar coatings of nano-needles facilitated the binding, stabilisation, and controlled release of BMP2 from the material which further enhanced MSC cell proliferation and bone formation. This study thus outlines a method for fabricating scaffolds with precise fibrous micro-architectures and bone mimetic nano-needle HA extrafibrillar coatings which significantly enhance MSC osteogenesis and therapeutic delivery and thus hold great promise for bone tissue regeneration.

## 7 Funding

The authors would like to acknowledge funding from European Research Council (ERC) Starting Grant (336882), Science Foundation Ireland (SFI) Support Grant SFI 13/ERC/L2864 and Irish Research Council Postgraduate Scholarship (GOIPG/2014/493).

